# Systematic engineering of plant cytochrome P450 system identifies a comprehensive strategy for expression of highly functional P450 enzymes in *Escherichia coli*

**DOI:** 10.1101/2022.12.13.520134

**Authors:** Michal Poborsky, Christoph Crocoll, Mohammed Saddik Motawie, Barbara Ann Halkier

## Abstract

Cytochrome P450s catalyse diverse and unique chemical reactions, which makes them invaluable enzymes in nature and industry. Metabolic engineers leverage these unique catalytic properties when refactoring plant biosynthetic pathways into microbial cell factories. However, due to their hydrophobic anchor, microbial expression of membrane-bound cytochrome P450s is challenging. An arsenal of protein engineering strategies was developed to improve their expression in *Escherichia coli*, but extensive screening is often necessary to tailor the engineering approach to an individual enzyme. Here, we propose a universal strategy that allows the expression of highly active cytochrome P450s in *E. coli* by systematically evaluating six common N-terminal modifications and their effect on *in vivo* activity of enzymes from the CYP79 and CYP83 families. We identified transmembrane domain truncation as the only strategy that had a significantly positive effect on all seven tested enzymes, increasing product titres between 2- to 170-fold. When comparing the changes in protein titre and product generation, we show that higher expression does not always translate to higher *in vivo* activity, thus making protein titre an unreliable screening target. Our results demonstrate that transmembrane domain truncation improves *in vivo* activity across a broad range of cytochrome P450s with diverse N-terminal sequences and could be applied as the modification-of-choice to avoid the time-consuming screening process and accelerate the future design of *E. coli* cell factories.

## INTRODUCTION

Cytochromes P450 (CYPs or P450s) are amongst the most versatile enzymes in nature and draw the attention of metabolic engineers with their ability to catalyse unique reactions on a myriad of complex substrates. *In planta*, P450s unlock the diversity of secondary metabolism (1) and play a key role in the biosynthesis of natural products used as medicines, cosmetics, colourants, and flavours (2, 3). Reconstituting these biosynthetic pathways in microbial cell factories, such as *Escherichia coli* or *Saccharomyces cerevisiae*, allows producing high-value natural products without the problems associated with the native hosts: seasonal nature of the production, low amounts of the product per plant, and tedious extraction from plant material. Notorious examples include the semi-synthetic production of the antimalarial drug artemisinin (4); biosynthesis of taxadiene, an intermediate of the anti-cancer medicine Taxol (5); and *de novo* biosynthesis of analgesic opioids (6, 7). Still, functional expression of heterologous P450s remains a challenge that researchers must address to reach competitive production levels of plant pathways in microbes.

All plant P450s require a redox partner for electron transfer and most localise on the membrane of the endoplasmic reticulum (8, 9), making *E. coli* - without eukaryotic organelles and no native P450 enzymes – a particularly challenging host. The N-terminal hydrophobic amino acids, which anchor P450s in membranes and act as a membrane localisation signal, cause the proteins to aggregate in insoluble inclusion bodies if the proteins cannot assume their native fold at a pace matching the speed of translation (10, 11). Larson et al. pioneered the strategy of N-terminal truncation, removing the hydrophobic amino acids of rabbit CYP2E1 and showing that the enzyme maintains its original activity (12). Barnes et al. optimised the 5’ codons of bovine CYP17A, increasing the translation initiation speed and enabling the protein to express in *E. coli* (13). N-terminal truncation and insertion of the MALLLAVF peptide, the so-called Barnes sequence, were later used to establish the expression of many other eukaryotic P450s in bacteria (14–18). Because these modifications can increase the ratio of P450s localised in the cytosol (19, 20), alternative techniques were developed to maintain P450 membrane localisation: insertion of leader sequences from bacterial membrane proteins (21) and exchange of the N-terminal domain with transmembrane sequences from *E. coli* membrane proteins (22, 23) or well-expressed eukaryotic P450 enzymes (24).

Although researchers can now choose from an arsenal of engineering strategies, no clear favourite exists and establishing novel plant pathways requires extensive optimisation and screening of the P450 enzyme expression and functionality (22, 23). Considering that most of the available research has focused on increasing the P450 protein titre, which does not always translate to higher *in vivo* activity (25), it is difficult to use previous results to guide the P450 engineering strategy in the context of cell factories. Furthermore, plant biosynthetic pathways often rely on multiple P450-mediated steps (1), while most studies in the field aim to overexpress only a single P450 enzyme at a time (26).

Here, we suggest a universal strategy to express highly functional P450s in *E. coli* cell factories by systematically comparing six common N-terminal modifications. We evaluate the changes of *in vivo* activity of CYP79A2 together with two members of the CYP83 family and then apply the best strategy to other P450 enzymes to test its universality. CYP79 and CYP83 catalyse two consecutive steps at the entrance to the glucosinolate pathway and accept standard amino acids as substrates (Fig. 1) (27). This makes them a convenient model system that avoids the need to feed complex substrates typical for plant secondary metabolism and allows the evaluation of concurrent engineering of multiple P450s.

**Fig. 1.**
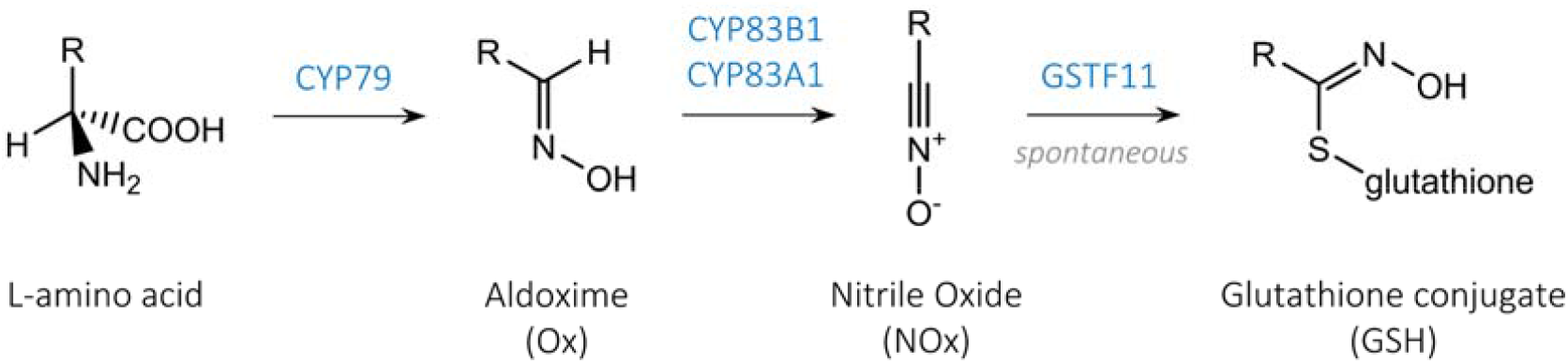
The first two steps of glucosinolate biosynthesis are catalysed by P450 enzymes from CYP79 and CYP83 families that convert amino acids into oximes and further into nitrile oxides, which can be conjugated with glutathione by a glutathione-S-transferase (GST) or spontaneously. We measured the generation of phenylalanine-derived products to evaluate the effect of N-terminal sequence modifications on P450 functionality in *E. coli:* phenylacetaldoxime (Phe-Ox), when engineering CYP79A2 alone, and *S*-phenylacetohydroxymoyl-L-glutathione (Phe-GSH), when engineering CYP79A2 together with CYP83s.

## RESULTS

### N-terminal sequence modifications affect CYP79A2 functionality and expression levels

We chose the conversion of phenylalanine to phenylacetaldoxime (Phe-Ox) catalysed by CYP79A2 for the initial assessment of N-terminal modifications because CYP79A2 constitutes a bottleneck in its biosynthetic pathways (27, 28), and it could be improved by N-terminal truncation (29) and the introduction of Barnes sequence (28). We cloned the full-length, codon-optimized CYP79A2 from *Arabidopsis thaliana* and six engineered variants with different N-terminal modifications into operons with A. *thaliana* reductase 1 ATR1 (Fig. 2D). All tested modifications were previously shown to increase the functional expression of P450s and fall into four general categories: [1] N-terminal truncation by removing either the transmembrane domain (ΔTM) or both the transmembrane domain and the adjacent hydrophilic region (ΔTM+) (12, 29), [2] 5’-codon optimisation by inserting short expression enhancing peptides in front of the P450 protein sequence (Barnes and 28aa), [3] transmembrane domain exchange with *E. coli* native membrane anchor from SohB probable protease (SohB) (22), and [4] introduction of *E. coli* leader sequence from outer membrane protein A (OmpA) (21) (Fig. 2A).

**Fig. 2.**
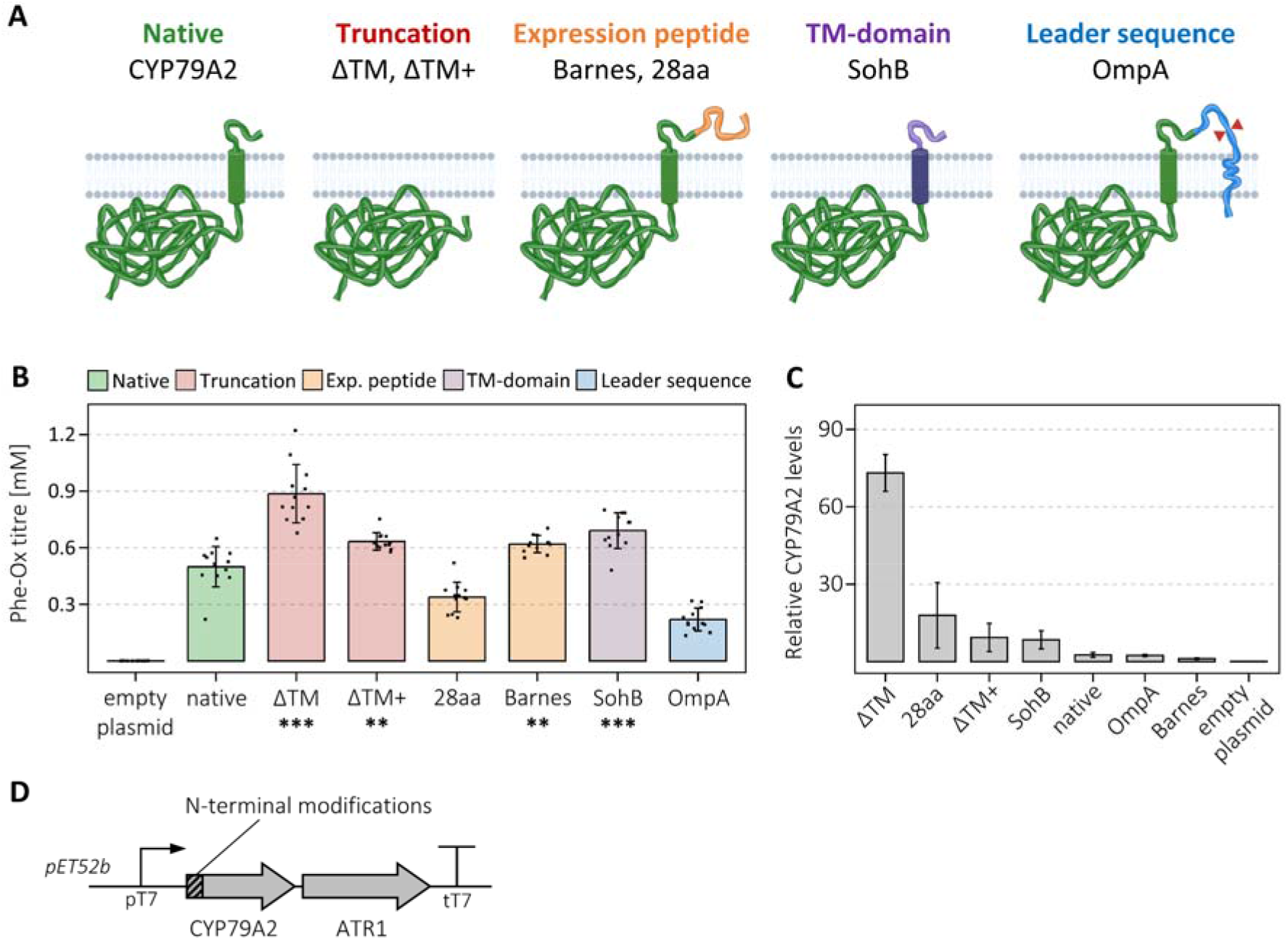
Effect of CYP79A2 N-terminal modifications on the production of phenylacetaldoxime (Phe-Ox). **A** N-terminal sequence modifications used to improve function of P450 enzymes include: (1) truncation of the transmembrane domain (ΔTM) or both transmembrane domain with adjacent hydrophilic region (ΔTM+), (2) expression enhancing peptides MALLLAVF(Barnes) or 28 amino acid tag (28aa), (3) transmembrane domain exchange with *E. coli* native probable protease (SohB), and (4) insertion of leader sequence from outer membrane protein A (OmpA) in front of full-length protein. **B** Phe-Ox titers produced from phenylalanine by native and engineered variants of CYP79A2. All strains were grown in 12 biological replicates across two independent experiments. Error bars represent standard deviation from the mean and outliers were removed from the data. Student’s upper-tailed *t* test denotes a significant increase in Phe-Ox titre compared to native with *p* value (with Bonferroni adjustment), * *p* < 0.05, ** *p* < 0.01, *** *p* < 0.001. **C** Relative quantification of CYP79A2 levels in strains expressing the native and engineered variants of CYP79A2, showing a representative peptide of CYP79A2 normalized to expression of *E. coli* isocitrate dehydrogenase. The bars represent the mean of 3-6 biological replicates from one of the experiments in B and error bars represent standard deviation from the mean. Additional CYP79A2 peptides can be found in Fig. S1. **D** Illustration of the CYP79A2-ATR1 operons as cloned into pET52b vector. Full-length cytochrome P450 oxidoreductase (ATR1) was co-expressed to supply electrons for the regeneration of P450 enzymes.

We assayed CYP79A2 variants by measuring the generation of Phe-Ox after 72 h fermentation in T7 autoinducing media (Fig. 2B). Although CYP79A2 was functional without any sequence modifications, producing 0.52 mM Phe-Ox, many engineered variants reached significantly higher titres. Those with the transmembrane domain removed or replaced were the top producers and [ΔTM]CYP79A2, where we truncated the transmembrane domain, but left the hydrophilic region intact, reached the highest Phe-Ox titre at 0.89 mM. The introduction of the Barnes sequence was beneficial only when the MALLLAVF peptide was inserted in front of the full-length enzyme, rather than substituting the first eight amino acids (Fig. S2). Using OmpA leader sequence or 28aa tag impaired CYP79A2 function, reducing Phe-Ox titre by approximately 30%. Because ATR2 and Δ44ATR2 are often used in literature, we compared the different reductases and their effect on [ΔTM] CYP79A2 activity. In our expression system, ATR1 was superior to both and led to 2-fold and 3-fold higher Phe-Ox titres than ATR2 and Δ44ATR2, respectively (Fig. S3) and was therefore used in all following experiments.

To relate the metabolite data with the P450 protein titre, we performed targeted proteomics measuring CYP79A2 protein expression levels on six of the replicates used above (Fig. 2C). We observed a disparity between the effect the modifications had on CYP79A2 protein expression compared to the increase in product generation. [ΔTM] CYP79A2 showed the highest protein levels and highest Phe-Ox production, but the relative protein titre increased by 28-fold, while Phe-Ox titre improved by only 1.8-fold. The introduction of 28aa tag increase protein levels 7-fold compared to native enzyme, while decreasing Phe-Ox production by 40%. These results corroborate previous reports that protein titre is not indicative of *in vivo* activity of P450 enzymes (25). In the following sections, we continue with [ΔTM]CYP79A2 as the best-performing variant using product generation as the metric to evaluate our engineering strategies.

### Concurrent engineering of a dual P450 module

Plant biosynthetic pathways for valuable secondary metabolites often proceed through multiple P450-mediated steps (1). To study the simultaneous engineering of two P450 enzymes, we paired native and truncated CYP79A2 with variants of CYP83B1 or CYP83A1, which both convert Phe-Ox into phenylacetonitrile oxide (Fig. 1). We expressed CYP79A2 with either one of the CYP83s and ATR1 from a single operon under the control of T7 promoter (Fig. 3A). When assaying the dual P450 module, we measured the generation of S-phenylacetohydroxymoyl-L-glutathione (Phe-GSH, Fig. 1), a more suitable product to quantify than the reactive and unstable nitrile oxide (30). Although the conjugation of nitrile oxides with glutathione can occur spontaneously, we co-expressed glutathione-S-transferase F11 (GSTF11) alongside the P450s and ATR1 to improve the formation of Phe-GSH (Fig. 3A) (31).

**Fig. 3.**
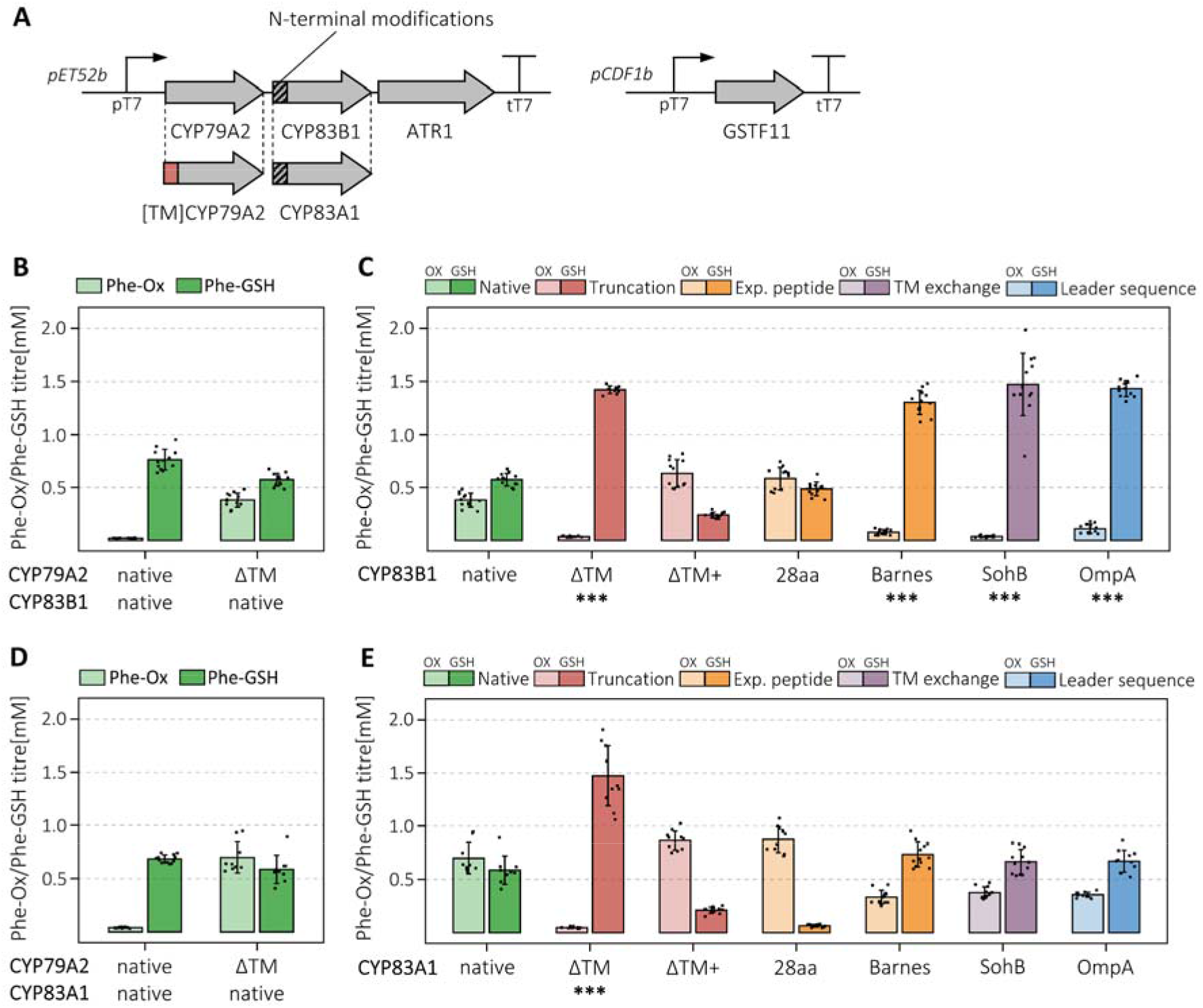
Engineering of the dual P450 module of CYP79A2 with CYP83B1 and CYP83A1. **A** The design of the expression constructs to examine the combinatorial effect of engineering the two P450s together. We paired the native and [ΔTM]-CYP79A2 with variants of the two CYP83s and co-expressed the P450 oxidoreductase ATR1 and glutathione transferase F11. **B, D** Production of phenylacetaldoxime (Phe-Ox, light colours) and *S-* phenylacetohydroxymoyl-L-glutathione (Phe-GSH, dark colours) by native and [ΔTM]CYP79A2 with native CYP83B1 and CYP83A1, respectively. **C, E** Production of Phe-Ox and Phe-GSH by [ΔTM]CYP79A2 with native and modified variants of CYP83B1 and CYP83A1, respectively. Bars are colour coded by the type of modification applied to CYP83 enzymes. All strains were grown in 12 biological replicates across two independent experiments. Error bars represent standard deviation from the mean and outliers were removed from the data. For **B-E**, Student’s upper-tailed *t* test denotes a significant increase in Phe-GSH titre compared to [ΔTM]CYP79A2 and native CYP83s with *p* value (with Holm adjustment), * *p* < 0.05, ** *p* < 0.01, *** *p* < 0.001.

We observed that strains with native CYP83s failed to convert the increased supply of Phe-Ox by [ΔTM] CYP79A2 into nitrile oxide, producing less Phe-GSH than with native CYP79A2 and accumulating Phe-Ox instead (Fig 3B, 3D). This was not specific to the truncated CYP79A2, as expressing the SohB variant led to the same result (Fig. S4). To achieve nearly complete conversion, we had to engineer the N-termini of both CYP79A2 and CYP83s. Although the highest Phe-GSH titre was similar between the CYP83s, interestingly, the N-terminal modifications’ effect on the two enzymes was different. For CYP83B1, multiple variants showed a significant increase in product titre with ΔTM, SohB and OmpA variants generating approximately 1.5 mM Phe-GSH, more than double compared to the native control (Fig. 3C). For CYP83A1, only ΔTM variant had a significantly positive effect on product titre and generated 1.47 mM Phe-GSH, on par with CYP83B1 (Fig. 3E). Our results suggest ΔTM N-terminal truncation is superior to the other approaches as it was the only modification that significantly increased the product generation by all three enzymes CYP79A2, CYP83B1 and CYP83A1.

### Transmembrane domain truncation as a universal strategy for P450 functional expression in *E. coli*

To validate transmembrane domain truncation as a strategy that could be applied to a broad range of P450s, we chose four additional CYP79 enzymes with diverse N-terminal sequences, distinct substrate preferences and different plant origins. CYP79A1 from *Sorghum bicolor* converts tyrosine into its corresponding oxime (32), CYP79B2 from A. *thaliana* accepts tryptophan as the sole substrate (33), CYP79D2 from *Manihot esculenta* acts on both valine and isoleucine (34), and CYP79F6 from *Barbarea vulgaris* accepts only non-canonical, carbon chain-elongated amino acids (here homophenylalanine) (35). We cloned the four CYP79s, CYP83 and ATR1 into operons

We assayed the enzymes by measuring the generation of putative glutathione conjugates derived from their respective amino acid substrates by tandem mass spectrometry (Q-TOF LC-MS/MS), comparing the relative production in strains expressing the native and truncated P450s (Fig. 4). We identified the glutathione conjugates by extracting molecular features with matching *m/z* ratios and a characteristic fragmentation pattern in the mass spectrum (36, 37), which we confirmed with Phe-GSH standard (Fig. S5). The results further corroborated the broad applicability of ΔTM as the transmembrane domain truncation led to higher product accumulation for all four new CYP79s. Although CYP79A1 and CYP79B2 were functional without any modification, truncating the enzymes yielded almost double of their respective products. The strain with [ΔTM]CYP79F6 and [ΔTM]CYP83A1 produced almost 7-fold more homophenylalanine-derived glutathione conjugate than the strain with native P450s and in the case of CYP79D2, which showed only trace activity without truncation, the improvement was 150-fold and 170-fold for valine and isoleucine products, respectively. Taken together, we show that transmembrane domain truncation improves functional expression and *in vivo* product accumulation for all seven studied P450 enzymes, being especially effective for enzymes with low native activity.

**Fig. 4.**
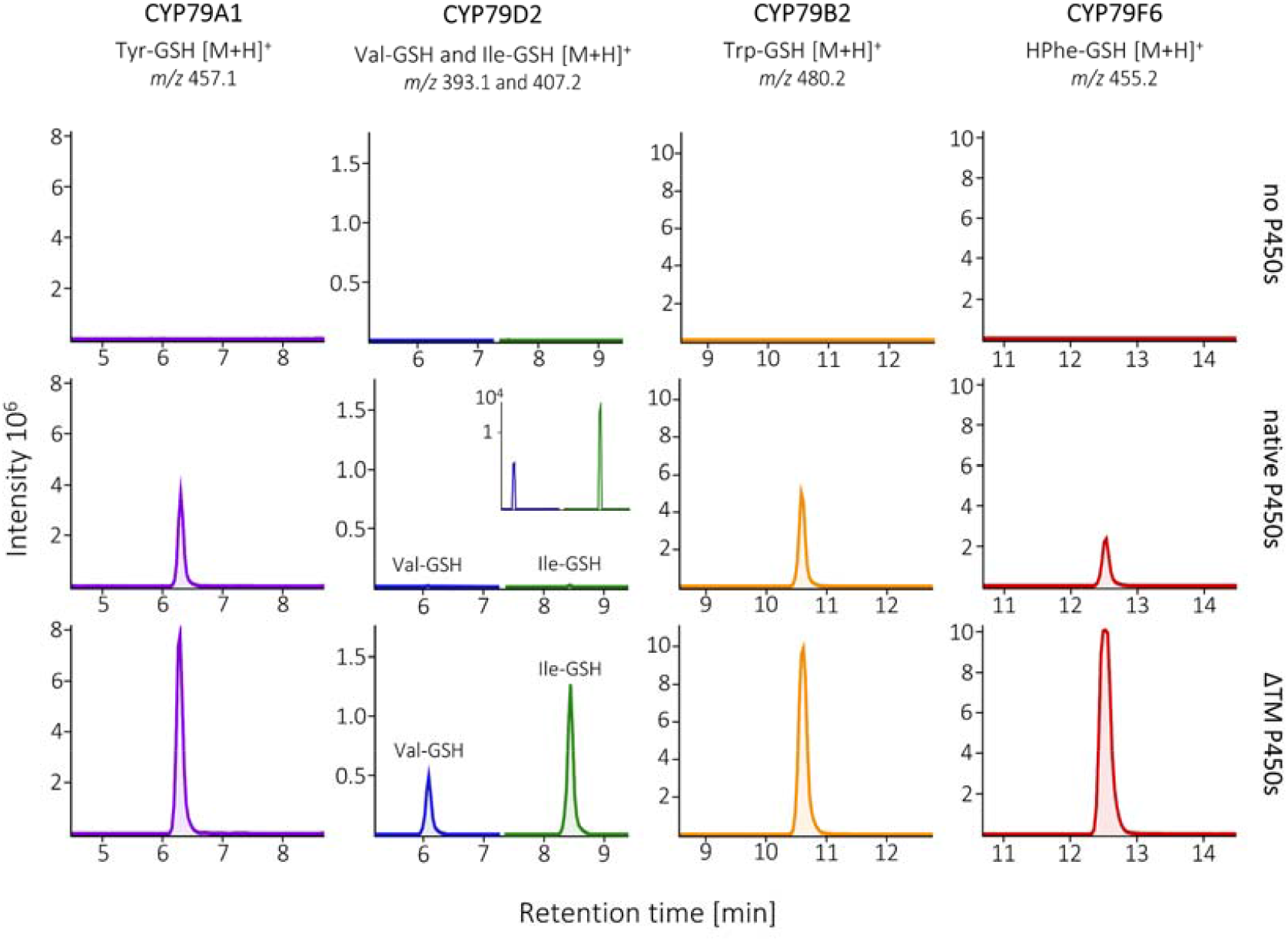
Validation of the transmembrane domain truncation (ΔTM) as a universal strategy on four more enzymes from CYP79 family with distinct substrate specificities and different plant origin. We expressed native and ΔTM variants of CYP79A1 (tyrosine-specific), CYP79D2 (valine- and isoleucine-specific), CYP79B2 (tryptophan-specific) and CYP79F6 (homophenylalanine (HPhe)-specific) from *Sorghum bicolor, Arabidopsis thaliana, Manihot esculenta*, and *Barbarea vulgaris* together with CYP83, ATR1 and GSTF11. We measured the generation of putative glutathione conjugates derived from the respective amino acids by Q-TOF LC-MS/MS. The presented extracted ion chromatograms are representative of 4 biological replicates and the characteristic fragmentation patters for each glutathione conjugate are shown in Fig S5. where both CYP79 and CYP83 were either native full-length or with the transmembrane domain truncated.

## DISCUSSION

We produced a broad and systematic screening of six common P450 N-terminal sequence modifications evaluating their effect on the *in vivo* enzymatic activity. Using two consecutive steps catalyzed by CYP79A2 and either CYP83A1 or CYP83B1 as a model system, we found that truncation of the transmembrane domain outperformed the other N-terminal modifications as it significantly improved product generation across the consecutive P450 enzymes. We sought to validate this finding by truncating four additional CYP79 enzymes and showed ΔTM variants led to increased product titres in all four cases. Amongst the six modifications, only the transmembrane domain truncation was beneficial for all studied combinations of enzymes from CYP79 and CYP83 families, improving the product titres 2- to 170-fold. This strongly suggests that the transmembrane domain truncation serves as a universal strategy to improve the functional expression of eukaryotic P450s in *E. coli.* Moreover, in the context of microbial cell factories, we demonstrated that in pathways with multiple P450s concentrating on the rate-limiting step is insufficient and all P450s must be engineered concurrently.

Classically, P450 engineering studies measure protein titre as a metric of success (25, 26), however, we observed a disparity between the effect the modifications had on the protein levels and the *in vivo* activity of CYP79A2 enzyme. ΔTM, ΔTM+ and SohB variants improved protein expression far greater than product titre and [28aa] CYP79A2 increased protein levels 7-fold while reducing Phe-Ox production by 30% compared to the native enzyme. Zhou et al. reviewed several studies that report a similar disconnection (25) and Christensen et al. showed that CYP79A1 fused to a membrane anchor of *E. coli* signal peptidase 1 was amongst the variants with the highest protein levels but only marginally improving *in vivo* activity (22). Together, these results indicate that screening P450 protein expression levels is insufficient to guide the engineering strategy. Screening P450 *in vivo* activity is more exact, however, it becomes cumbersome at scale because it requires access to diverse and complex substrates as well as established quantification methods for the generated products.

To circumvent the screening process, we set out to find a broadly applicable engineering strategy, as, in our opinion, no clear favourite has emerged in literature. Available studies use different growth conditions and combine multiple modifications to tailor individual P450 enzymes for the highest possible protein titre, making it difficult to abstract a general engineering strategy. For example, N-terminal truncations were previously successful in enhancing the expression of many eukaryotic P450s in bacteria, supporting our findings, but the truncations varied in length and were often complemented by Barnes sequence (17, 32), optimized translation initiation region (10, 38), or amino acid substitutions downstream in the protein sequence (16, 39). We studied the common modifications individually and found that only ΔTM caused a significant increase in product generation across all tested enzymes, and while other modifications led to similar product titres in certain cases, they failed to have a consistently positive effect. Barnes and SohB variants were significantly better than native CYP79A2 and CYP83B1 but caused no statistically significant change in CYP83A1. OmpA was among the top variants for CYP83B1 but led to a dramatic decrease in Phe-Ox production by CYP79A2. ΔTM+ was successful only in the case of CYP79A2 as both CYP83 ΔTM+ variants performed poorly. These results suggest the transmembrane domain as the major factor that impedes the functional expression of P450s in *E. coli*, and its truncation is sufficient to alleviate the problem in a broad range of P450 enzymes.

Balancing the expression of individual enzymes or modules is a staple of pathway engineering. We noticed a similar requirement for P450 engineering when targeting a single rate-liming enzyme out of multiple P450s in a pathway was not enough to resolve the bottleneck and only moved it further downstream. We showed that [ΔTM]CYP79A2, which catalyses the rate-limiting reaction at the entry point of the glucosinolate pathway (27, 28), produced almost double Phe-Ox over the native enzyme, but this benefit was lost upon adding the subsequent unmodified CYP83s, which were unable to convert the extra Phe-Ox into nitrile oxide. Only when engineering both CYP79A2 and CYP83s, we successfully translated the doubled Phe-Ox titre into doubled Phe-GSH. As P450s draw from shared cellular resources such as heme incorporation, electron supply through NADPH, and electron transfer from NADPH to P450 by cytochrome P450 reductase, the 28-fold increase in expression we observed for [ΔTM]CYP79A2 might cause CYP79A2 to monopolize the access to these resources and, in turn, impair CYP83 activity.

In conclusion, we found that transmembrane domain truncation universally improves the functionality of P450 enzymes in *E. coli.* Our results emphasise the disparity between P450 *in vivo* activity and protein titre and suggest that screening product generation is necessary to guide P450 engineering. To avoid extensive screening, we propose ΔTM as a ‘modification of choice’ that yields highly functional P450 enzymes and aids the future design and establishment of P450-containing biosynthetic pathways in *E. coli* cell factories.

## MATERIALS AND METHODS

### Oligonucleotides, genes, plasmids, and strains

All cloning work was performed in *E. coli* NEB 10-β and BL21 (DE3) strain (both from New England BioLabs, Ipswich, USA) with pRI952 plasmid used for all fermentations. pRI952 plasmid carries genes for rare tRNAs for isoleucine and arginine (40). All plasmids used in this study can be found in Supplementary table 2. Plant genes were ordered codon optimized from either Integrated DNA Technologies (IDT, Leuven, Belgium) or Twist Bioscience (San Francisco, CA, USA). Oligonucleotides were purchased from IDT (Leuven, Belgium) and Sanger sequencing was provided by Eurofins Genomics (Ebersberg, Germany). Omega Bio’s bacterial kits were used for all DNA preparation.

### DNA assembly

Proofreading Phusion U Hot Start DNA polymerase (Thermo Fisher Scientific, Waltham, MA) with uracil-containing oligonucleotides was used to amplify all DNA parts. pET-52b (Novagen®, Merck, #71554) and pCDF-1b (Novagen®, Merck, #71330) plasmids were used to express the heterologous genes. To assemble purified PCR products, the USER protocol was adapted from Geu-Flores et al. (41) and Cavaleiro et al. (42) in a 10 μL reaction as follows: 1 μL USER enzyme mix, 1 μL 10x CutSmart buffer, 1 unit of DpnI, 20 ng PCR product of the plasmid backbone and 40 ng PCR product of all fragments to assemble. All components of USER assembly were purchased from New England BioLabs, Ipswich, USA. The reaction was incubated for 1 h at 37 °C, then at gradually decreasing temperatures around the melting temperature (T_m_) of USER overhangs (31-26 °C) with 5 min at each temperature step and finished with 30 min at 10 °C. Afterwards, the entire reaction was transformed into chemically competent *E. coli* NEB 10-β cells. Colony PCR was performed with DreamTaq DNA polymerase (Thermo Scientific, Waltham, USA), and the sequences of cloned constructs were verified with Mix2Seq kits (Eurofins Genomics, Germany).

### Construction of P450 expression modules with N-terminal modifications

N-terminal modifications of CYP79 and CYP83 sequences included: (1) truncation of the membrane domain in front of the hydrophilic region (ΔTM), (2) truncation of the membrane domain after the hydrophilic region (ΔTM+), (3) addition of Barnes sequence MALLLAVF in front of the full-length P450 sequence (Barnes), (4) substitution of the first 8 amino acids with Barnes sequence MALLLAVF, (5) addition of 28 amino acid synthetic peptide in front of the full-length P450 sequence (28aa), (6) substitution of the native transmembrane domain with the transmembrane domain of SohB probable protease from *E. coli* (SohB) and (7) addition of *E. coli* outer membrane protein A (OmpA) signal peptide in front of the full-length P450 sequence (OmpA). 28aa, SohB and OmpA sequences were ordered from Twist Bioscience as short DNA fragments, amplified with uracil-containing primers, and used to create modified P450s on a separate plasmid backbone before they were cloned into the expression vector. Barnes sequence was short enough to be inserted into the primer overhangs that were used to create the modifications (3) and (4), and truncated P450s were prepared with primers annealing at within the gene sequences. DNA sequences of all used modifications are shown in Supplementary table 1. When expressing multiple genes from a single plasmid, the genes were assembled into synthetic operons with a maximum of three genes per operon. All operons were controlled by consensus T7 promoters with T7 terminators, and all genes shared the same ribosome binding site originally from the T7 bacteriophage major capsid protein. To define transmembrane domains of P450 enzymes, we used uniport annotations in combination with a transmembrane domain prediction tool TMHMM v2.0 (now depreciated in favour of DeepTHMHH) (43) (Table 1). To identify an appropriate truncation site for ΔTM+, the hydrophilic region was annotated as a short amino acid stretch abundant in arginine and lysine residues together with other hydrophilic amino acids.

**Tab. 1.** Seven P450 enzymes of the glucosinolate pathway from four different plants were targeted for engineering in our study. CYP79s control the amino acid substrate entering the pathway and we selected five with distinct substrate preferences and plant origin. The subsequent CYP83s are promiscuous and convert any oximes generated by CYP79s. ΔTM and ΔTM + represent the residues that were truncated after the membrane anchor or after the membrane anchor with the hydrophilic region.

| Enzyme | Substrates | ΔTM | ΔTM+ | Plant origin |
| --- | --- | --- | --- | --- |
| CYP79A1 | Tyr | Δ38 | - | *Sorghum bicolor* |
| CYP79A2 | Phe | Δ14 | Δ23 | *Arabidopsis thaliana* |
| CYP79B2 | Trp | Δ42 | - | *Arabidopsis thaliana* |
| CYP79D2 | Val, Ile | Δ41 | - | *Manihot esculenta* |
| CYP79F6 | HPhe | Δ29 | | *Barbarea Vulgaris* |
| CYP83A1 | AA oximes | Δ23 | Δ28 | *Arabidopsis thaliana* |
| CYP83B1 | AA oximes | Δ21 | Δ29 | *Arabidopsis thaliana* |

### Transformation of the expression strain

Allowing the transformation of up to 4 plasmids at the same time, electrocompetent cells were always prepared fresh rather than using frozen aliquots. 30 μL of overnight culture or a single colony were inoculated into 1.4 mL LB media in a microcentrifuge tube and grown for 3-4 hours at 37 °C, 900 RPM in a shaking heating block. Afterwards, the cells were washed twice with 1 mL of ice-cold water and transformed with 1 μL of each plasmid. The GenePulser Xcell (Bio-Rad, Hercules, USA) electroporator was set to 1350 V, 10 μF, 600 Ohms. The transformants were recovered in SOC medium for 1 h at 37 °C, 900 RPM, spread on LB agar with appropriate antibiotics and grown overnight at 37 °C. Before fermentation, single colonies were picked into 1.2 mL LB media with appropriate antibiotics in a 24-well plate (Thermo Fisher Scientific, Waltham, MA) and the precultures were cultivated overnight.

### Induction of protein expression and fermentation conditions

Autoinducing medium, adapted from Studier et al. (44), was used during all fermentations unless stated otherwise. The original recipe was adjusted to resemble Terrific broth (TB) in the yeast extract, tryptone and buffer composition, as TB is a common choice when expressing P450s. The TB-based autoinducing medium contained 24 g/L yeast extract, 12 g/L tryptone, 100 mM KH_2_PO_4_-K_2_HPO_4_ buffer pH 7.0, 0.05% glucose, 0.5% glycerol, 0.2% α-lactose, 2 mM MgSO_4_, and trace metal mix (50 μM FeCl_3_, 20 μM CaCl_2_, 10 μM MnCl_2_, 10 μM ZnSO_4_, 2 μM CoCl_2_, 2 μM CuCl_2_, 2 μM NiCl_2_, 2 μM Na_2_MoO_4_, 2 μM Na_2_SeO_3_, 2 μM H_3_BO_3_). To improve heme biosynthesis, which is necessary for functional P450 enzymes, 0.5 mM 5-aminolevulinic acid was supplied to the media. Required antibiotics were also added at the following concentrations: carbenicillin (50 μg/mL), spectinomycin (50 μg/mL), kanamycin (50 μg/mL) and chloramphenicol (34 μg/mL). Precultures were diluted 1000-fold into 1.2 mL autoinducing medium in round bottom 24-well plates and the fermentation was carried out in a rotary shaker at 18 °C, 200 RPM for 72 h.

### Liquid chromatography coupled to mass spectrometry (LC-MS) metabolite analysis

#### Harvesting E. coli media samples

Media samples were collected after cells were spun down from 1 mL cultures in deep 96-well plates at 3700 x *g* for 5 min. The collected samples were stored at −20 °C until they were analyzed by high-performance liquid chromatography tandem mass spectrometry (LC-MS/MS). Before analysis, an aliquot of the supernatant was diluted 25-fold in water and filtered through 0.22 μm filters.

#### Combined analysis of amino acids, phenylacetaldoxime and GSH-conjugates by LC coupled to triple quadrupole MS (LC-MS/QqQ)

The pre-diluted media samples were diluted further in a ratio of 1:10 (v:v) in water containing the ^13^C-, ^15^N-labeled amino acid mix (Isotec, Miamisburg, US). Amino acids in the diluted extracts were directly analyzed by LC-MS/MS. The analysis method originates from a protocol described by Jander et al. (45) and was modified from the protocol described in Docimo et al. (46) to adjust for instrument settings and inclusion of metabolites from the benzyl-glucosinolate pathway. Chromatography was performed on an Advance ultra-high-performance liquid chromatography (UHPLC) system (Bruker, Bremen, Germany). Separation was achieved on a Zorbax Eclipse XDB-C18 column (50 x 4.6mm, 1.8μm, Agilent Technologies, Germany). Formic acid (0.05%) in water (v/v) and acetonitrile (supplied with 0.05% formic acid, v/v) were employed as mobile phases A and B, respectively. The elution profile was: 0-1.2 min, 3 % B; 1.2-3.8 min, 3-65 % B; 3.8-4.0 min 65-100% B, 4.0-4.6 min 100% B, 4.6-4.7 min 100-3% B and 4.7-6.0 min in 3% B. The mobile phase flow rate was 500 μl/min. The column temperature was maintained at 40 °C. The liquid chromatography was coupled to an EVOQ Elite Triplequadrupole (QqQ) mass spectrometer (Bruker, Bremen, Germany) equipped with an electrospray ion source (ESI) operated in combined positive and negative ionization modes. The instrument parameters were optimized by infusion experiments with pure standards. The ion spray voltage was maintained at +3000 V or - 4000V for amino acid and glucosinolate analysis, respectively. The cone temperature was set to 300 °C and the cone gas to 20 psi. The heated probe temperature was set to 300 °C and the probe gas flow to 50 psi. Nebulizing gas was set to 60 psi and collision gas to 1.6 mTorr. Nitrogen was used as the probe and nebulizing gas and argon as the collision gas. The active exhaust was constantly on. Multiple reaction monitoring (MRM) was used to monitor analyte parent ion ⍰ product ion transitions: MRMs were chosen (28, 45–47)

Bruker MS Workstation software (Version 8.2.1, Bruker, Bremen, Germany) was used for data acquisition and processing. Linearity in ionization efficiencies was verified by analysing dilution series of standard mixtures (amino acid standard mix, Fluka plus Gln and Trp, also Fluka). All samples were spiked with ^13^C-, ^15^N-labeled amino acids at a concentration of 10 μg/mL. The concentration of the individual labelled amino acids in the mix had been determined by comparison to a reference standard by LC-MS/MS analysis. Individual amino acids in the sample were quantified by the respective ^13^C-, ^15^N-labeled amino acid internal standards, except for tryptophan and asparagine: tryptophan was quantified using ^13^C-, ^15^N-Phe applying a response factor of 0.42, asparagine was quantified using ^13^C-, ^15^N-Asp applying a response factor of 1.0. Concentrations of the produced Phe-Ox and Phe-GSH were quantified with ^13^C-, ^15^N-Phe as internal standard using response factors calculated from dilution series of the analytes in spent *E. coli* media to account for the matrix effect. Outliers were defined as Q1 – 2 * IQR, Q3 + 2 * IQR (Q = quartile, IQR = interquartile range) and removed from the data. Student’s upper-tailed *t* test was performed to identify engineered variants with significant increase in product generation compared to native enzymes. Statistical analysis and visualisation were done using R (version 4.2.2, (48)) in RStudio 2022.07.2.576 (49) relying on tidyverse (50) and ggplot2 (51) packages.

#### Untargeted analysis of glutathione conjugates by Q-TOF LC-MS/MS

Samples for Q-TOF analysis were prepared similarly to the triple quad, but at higher concentrations, only diluting the media 25-fold in water spiked with 1 μM caffeine as internal standard. To identify potential intermediates not included in the targeted identification by LC-MS/QqQ as described above, samples were also subjected to untargeted analysis by LC-MS/Q-TOF. Chromatography was performed on a Dionex UltiMate® 3000 Quaternary Rapid Separation UHPLC^+^ focused system (Thermo Fisher Scientific, Germering, Germany). Separation was achieved on a Kinetex 1.7u XB-C18 column (100 x 2.1 mm, 1.7 μm, 100 Å, Phenomenex, Torrance, CA, USA). Formic acid (0.05%) in water and acetonitrile (supplied with 0.05% formic acid) were employed as mobile phases A and B, respectively. The elution profile was: 0.00-0.1 min, 2% B; 0.1.-16.0 min, 2-45% B; 16.0-24.5 min 45-100% B, 24.5-26.5 min 100% B, 26.5-26.55 min 100-2% B and 26.55-30.0 2% B. The mobile phase flow rate was 300 μl/min. The column temperature was maintained at 25 °C. The liquid chromatography was coupled to a Compact micrOTOF-Q mass spectrometer (Bruker, Bremen, Germany) equipped with an ESI operated in positive mode. Settings for positive ion mode were as follows: the ion spray voltage was maintained at +4500 V. Dry temperature was set to 250 °C and dry gas flow was set to 8 L/min. Nebulizing gas was set to 2.5 bar and collision energy to 10 eV. Nitrogen was used as dry gas, nebulizing gas, and collision gas. Sodium formate (Na-formate) clusters were used as calibrant AutoMSMS mode employed to collect MS and MS/MS spectra of the three most abundant ions present. The acquisition rate was at 2 Hz. The *m/z* range was set to 50-1000 for MS acquisition and *m/z* 200-800 for MS/MS acquisition. All files were automatically calibrated based on the compound spectra collected from the Na-formate clusters by post-processing. The characteristic fragmentation pattern of GSH conjugates is described previously (37, 52) (Supplementary figure 5).

#### Targeted proteomics

Proteomic analysis method was adapted from Bathh et al. (53) as previously described (28).

## ACKNOWLEDGEMENTS

This work was supported by Danish National Research Foundation grant DNRF99 awarded to B.A.H. and Novo Nordisk Foundation grant NNF20OC0065061 awarded to B.A.H.

