## Supplementary data for "Systematic engineering of plant cytochrome P450 system identifies a comprehensive strategy for expression of highly functional P450 enzymes in *Escherichia coli*"

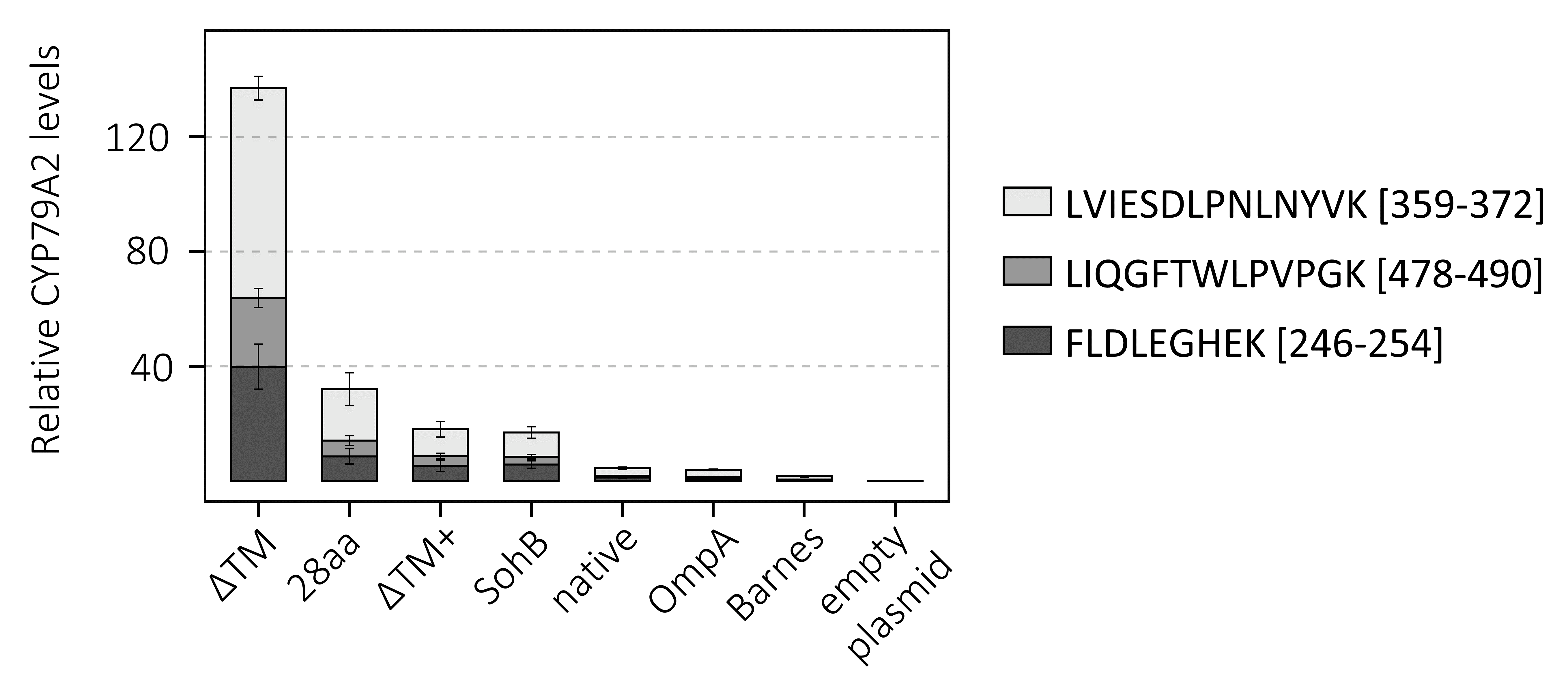

**Fig. S1** | Expression levels of CYP79A2 represented by three proteotypic peptides normalised to the expression of E. coli isocitrate dehydrogenase. The bars represent the mean of 6 biological replicates from of the experiments and error bars represent standard error of the mean.

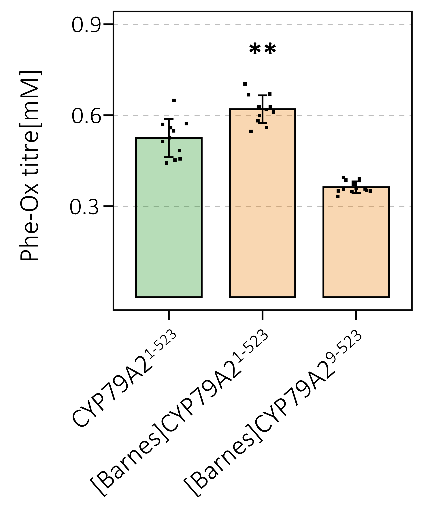

**Fig. S2** | Insertion of MALLLAVF peptide in front of full-length CYP79A2 or by substituting the first eight amino acids. Every strain was grown in 12 biological replicates and the error bars represent standard deviation from the mean. Student’s two-tailed *t* test notes significant increase in Phe-Ox titre compared to native with *p* value (with Holm adjustment), * *p* < 0.05, ** *p* < 0.01, *** *p* < 0.001.

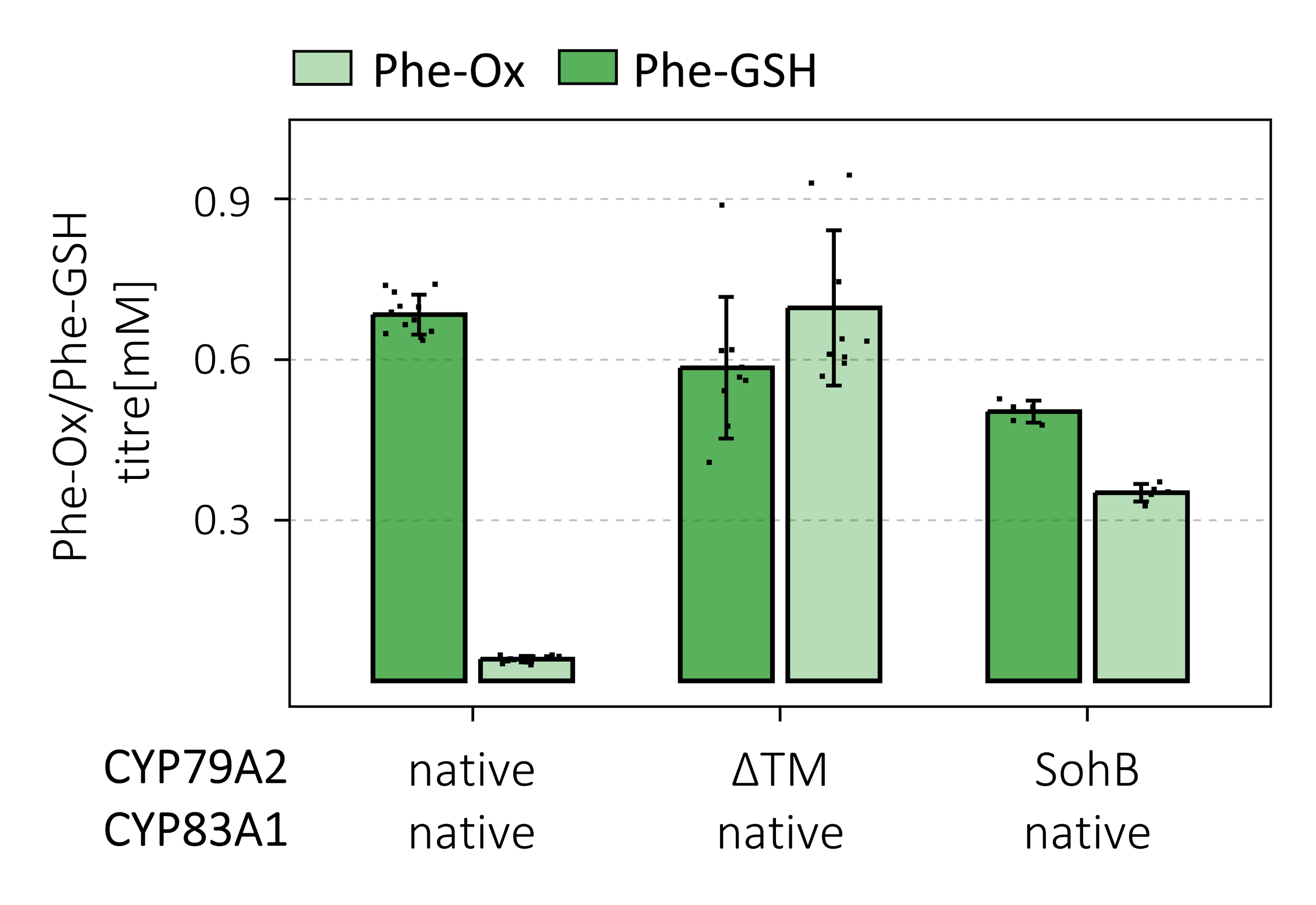

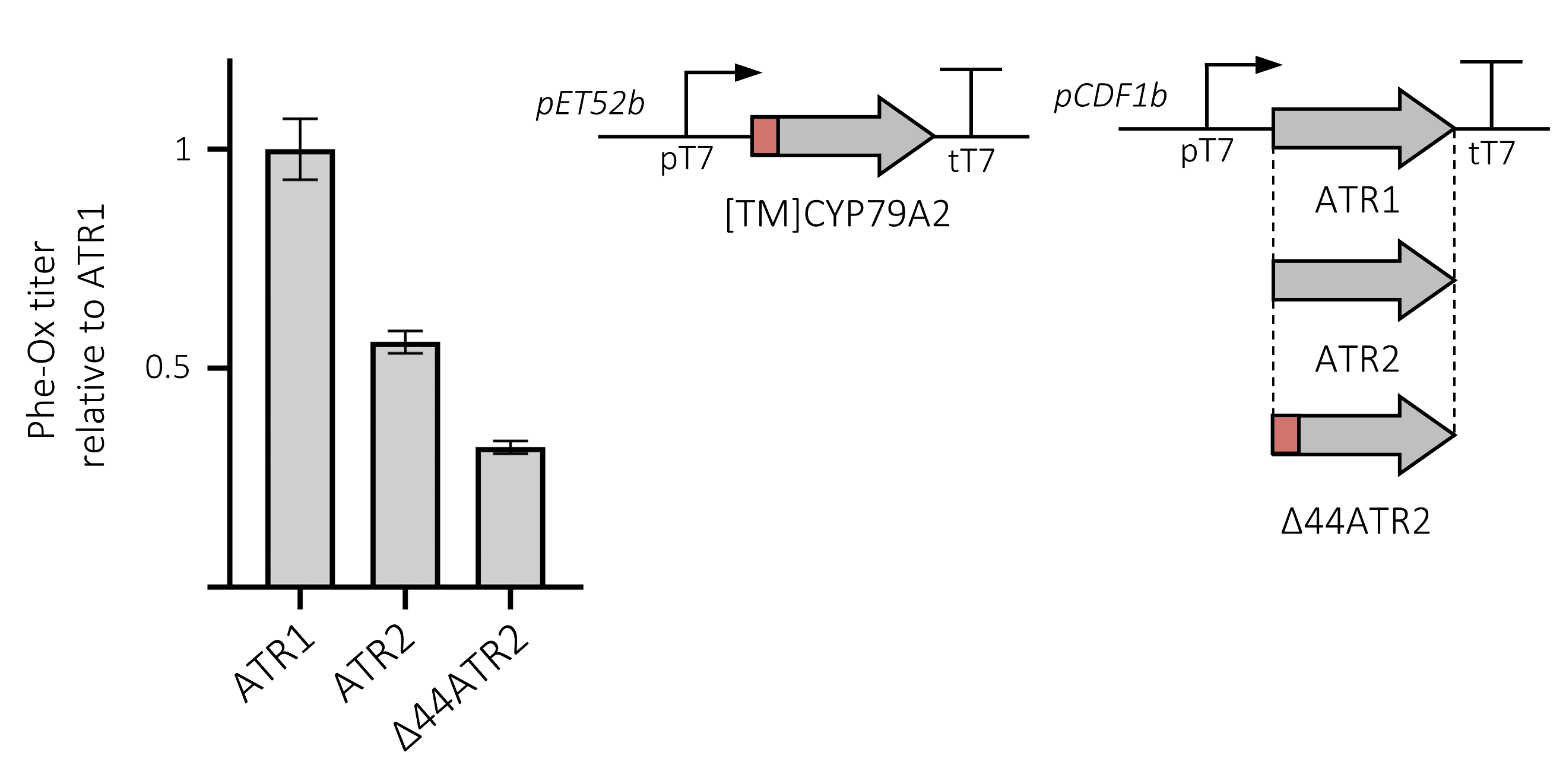

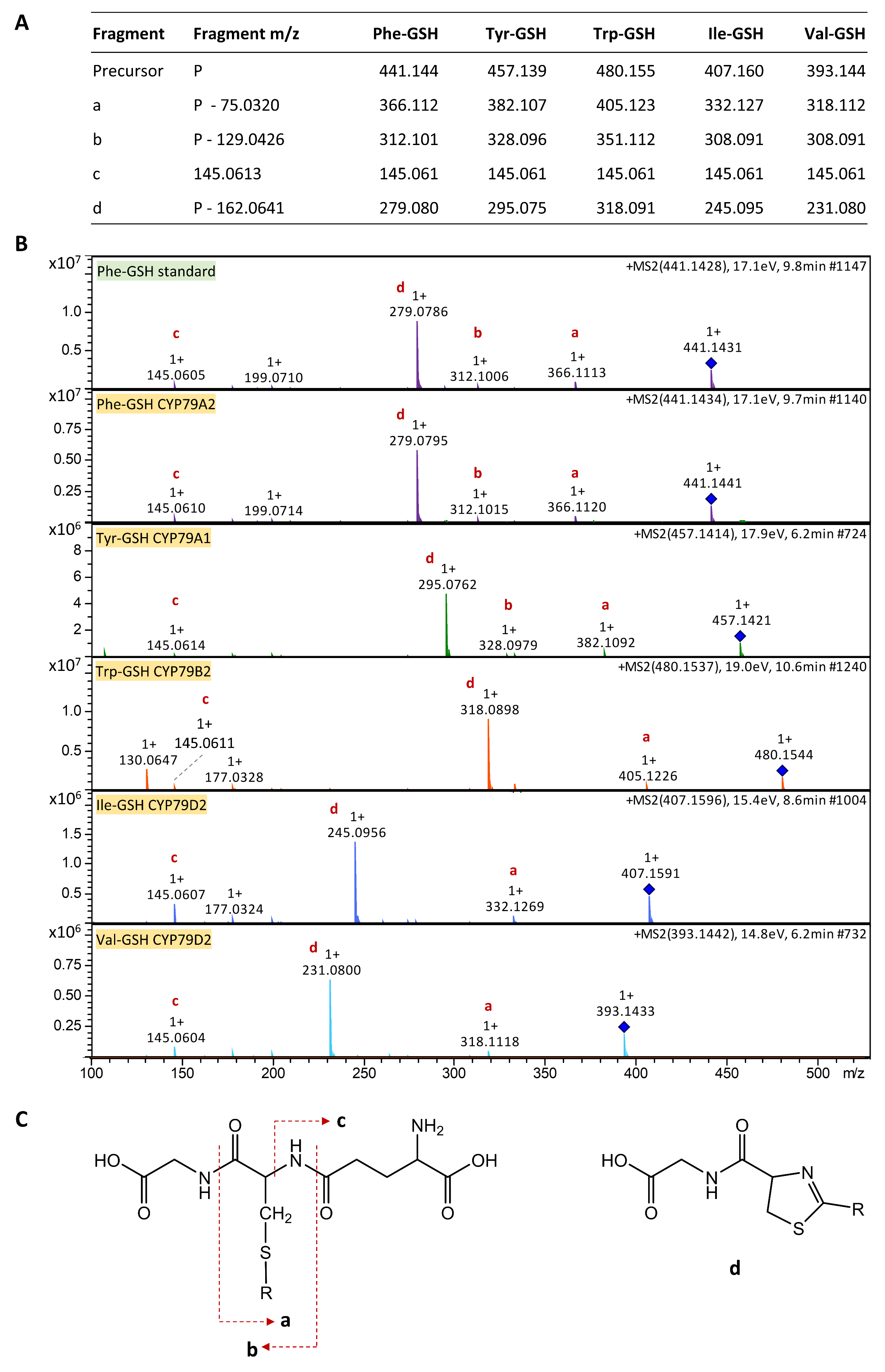

**Fig. S4** | Comparison of pairing native CYP83A1 with 2 different engineered variants of CYP79A2 in a similar setup as Fig 3D in the main text. Every strain was grown in 6 (SohB) or 12 (native and ΔTM) biological replicates and the error bars represent standard deviation from the mean.

**Fig. S3** | Testing of the two available cytochrome P450 reductases from *A. thaliana*, ATR1, ATR2 and its truncated variant Δ44ATR2. The protein expression was induced by addition of 0.5 mM Isopropyl ß-D-1-thiogalactopyranoside (IPTG) after the cultures reached OD600 of 0.6. Every strain was grown in 3 biological replicates and the error bars represent standard deviation from the mean.

**Fig. S5** | **A** Glutathione-conjugate characteristic fragments (refs) with masses corresponding to intermediates derived from specific amino acids. **B** The fragments are demonstrated using an authentic standard of GSH-conjugate with phenylacetonitrile oxide, an intermediate of benzyl glucosinolate biosynthesis, and then our samples upon expression of different CYP79s. **C** Molecular fragments of glutathione conjugates detected in our experiments.

**Tab. S1 |** Nucleotide sequences of all P450 modifications used in the study other than truncation.

| **Modification** | **Sequence 5'-> 3'** |
| --- | --- |
| Barnes | ATGGCTCTGTTATTAGCAGTTTTT |
| 28aa | ATGGAATTATCACAAGTTTGTACAAAAAAGGAGGCTGGCGCCGGAACCAATTCAGTCGACTGGATCCAAGAAGGAGATATAACC |
| SohB | ATGGAATTGTTGTCTGAATATGGTTTGTTTTTGGCGAAAATCGTTACCGTTGTGCTAGCGATTGCGGCGATTGCCGCCATTATTGTCAATGTTGCTCAACGTAATAAACGCCAGCGTGGCGAGTTACGGGTAAACAATCTTAGC |
| OmpA | ATGAAAAAGACAGCTATCGCGATTGCAGTGGCACTGGCTGGTTTCGCTACCGTAGCTCAGGCC |

**Tab. S2 |** List of all plasmids created and used during this study.

| **#** | **Plasmid** | **Description** | **Reference** |
| --- | --- | --- | --- |
| 1 | pCDF1b | used to construct expression vectors | Novagen |
| 2 | pET52b(+) | used to construct expression vectors | Novagen |
| 3 | pRI952 | carries ileX and argU | Del Tito et al., 1995 |
| 4 | pET52-CYP79A2-ATR1 | expression of CYP79A2-ATR1 operon | This study |
| 5 | pET52-Δ14CYP79A2-ATR1 | expression of [ΔTM]CYP79A2-ATR1 operon | This study |
| 6 | pET52-Δ23CYP79A2-ATR1 | expression of [ΔTM+]CYP79A2-ATR1 operon | This study |
| 7 | pET52-bCYP79A2-ATR1 | expression of [÷Barnes]CYP79A2-ATR1 operon | This study |
| 8 | pET52-Barnes-CYP79A2-ATR1 | expression of [+Barnes]CYP79A2-ATR1 operon | This study |
| 9 | pET52-[OmpA]CYP79A2-ATR1 | expression of [OmpA]CYP79A2-ATR1 operon | This study |
| 10 | pET52-[SohB]Δ14CYP79A2-ATR1 | expression of [SohB]CYP79A2-ATR1 operon | This study |
| 11 | pET52-[28aa]CYP79A2-ATR1 | expression of [28aa]CYP79A2-ATR1 operon | This study |
| 12 | pET52-CYP79A2-CYP83B1-ATR1 | expression of CYP79A2-CYP83B1-ATR1 operon | This study |
| 13 | pET52-Δ14CYP79A2-CYP83B1-ATR1 | expression of [ΔTM]CYP79A2-CYP83B1-ATR1 operon | This study |
| 14 | pET52-Δ14CYP79A2-Δ23CYP83B1-ATR1 | expression of [ΔTM]CYP79A2-[ΔTM]CYP83B1-ATR1 operon | This study |
| 15 | pET52-Δ14CYP79A2-Δ28CYP83B1-ATR1 | expression of [ΔTM]CYP79A2-[ΔTM+]CYP83B1-ATR1 operon | This study |
| 16 | pET52-Δ14CYP79A2-bCYP83B1-ATR1 | expression of [ΔTM]CYP79A2-[÷Barnes]CYP83B1-ATR1 operon | This study |
| 17 | pET52-Δ14CYP79A2-[Barnes]CYP83B1-ATR1 | expression of [ΔTM]CYP79A2-[+Barnes]CYP83B1-ATR1 operon | This study |
| 18 | pET52-Δ14CYP79A2-[OmpA]CYP83B1-ATR1 | expression of [ΔTM]CYP79A2-[OmpA]CYP83B1-ATR1 operon | This study |
| 19 | pET52-Δ14CYP79A2-[SohB]Δ23CYP83B1-ATR1 | expression of [ΔTM]CYP79A2-[SohB]CYP83B1-ATR1 operon | This study |
| 20 | pET52-Δ14CYP79A2-[28aa]CYP83B1-ATR1 | expression of [ΔTM]CYP79A2-[28aa]CYP83B1-ATR1 operon | This study |
| 21 | pET52-CYP79A2-CYP83A1-ATR1 | expression of CYP79A2-CYP83A1-ATR1 operon | This study |
| 22 | pET52-Δ14CYP79A2-CYP83A1-ATR1 | expression of [ΔTM]CYP79A2-CYP83A1-ATR1 operon | This study |
| 23 | pET52-Δ14CYP79A2-Δ21CYP83A1-ATR1 | expression of [ΔTM]CYP79A2-[ΔTM]CYP83A1-ATR1 operon | This study |
| 24 | pET52-Δ14CYP79A2-Δ29CYP83A1-ATR1 | expression of [ΔTM]CYP79A2-[ΔTM+]CYP83A1-ATR1 operon | This study |
| 25 | pET52-Δ14CYP79A2-bCYP83A1-ATR1 | expression of [ΔTM]CYP79A2-[÷Barnes]CYP83A1-ATR1 operon | This study |
| 26 | pET52-Δ14CYP79A2-[Barnes]CYP83A1-ATR1 | expression of [ΔTM]CYP79A2-[+Barnes]CYP83A1-ATR1 operon | This study |
| 27 | pET52-Δ14CYP79A2-[OmpA]CYP83A1-ATR1 | expression of [ΔTM]CYP79A2-[OmpA]CYP83A1-ATR1 operon | This study |
| 28 | pET52-Δ14CYP79A2-[SohB]Δ21CYP83A1-ATR1 | expression of [ΔTM]CYP79A2-[SohB]CYP83A1-ATR1 operon | This study |
| 29 | pET52-Δ14CYP79A2-[28aa]CYP83A1-ATR1 | expression of [ΔTM]CYP79A2-[28aa]CYP83A1-ATR1 operon | This study |
| 30 | pET52-[SohB]Δ14CYP79A2-CYP83A1-ATR1 | expression of [SohB]CYP79A2-CYP83B1-ATR1 operon | This study |
| 31 | pCDF-GSTF11 | expression of GSTF11 | This study |
| 32 | pET52-CYP79A1-CYP83B1-ATR1 | expression of CYP79A1-CYP83B1-ATR1 operon | This study |
| 33 | pET52-CYP79B2-CYP83B1-ATR1 | expression of CYP79B2-CYP83B1-ATR1 operon | This study |
| 34 | pET52-CYP79D2-CYP83B1-ATR1 | expression of CYP79D2-CYP83B1-ATR1 operon | This study |
| 35 | pET52-CYP79F6-CYP83A1-ATR1 | expression of CYP79F6-CYP83A1-ATR1 operon | This study |
| 36 | pET52-Δ38CYP79A1-Δ23CYP83B1-ATR1 | expression of [ΔTM]CYP79A1-[ΔTM]CYP83B1-ATR1 operon | This study |
| 37 | pET52-Δ42CYP79B2-Δ23CYP83B1-ATR1 | expression of [ΔTM]CYP79B2-[ΔTM]CYP83B1-ATR1 operon | This study |
| 38 | pET52-Δ41CYP79D2-Δ23CYP83B1-ATR1 | expression of [ΔTM]CYP79D2-[ΔTM]CYP83B1-ATR1 operon | This study |
| 39 | pET52-Δ29CYP79F6-Δ21CYP83A1-ATR1 | expression of [ΔTM]CYP79F6-[ΔTM]CYP83A1-ATR1 operon | This study |
| 40 | pCDF-ATR1 | expression of ATR1 | This study |
| 41 | pCDF-ATR2 | expression of ATR2 | This study |
| 42 | pCDF-Δ44ATR2 | expression of Δ44ATR2 | This study |
